# KLF2 is a therapeutic target for COVID-19 induced endothelial dysfunction

**DOI:** 10.1101/2021.02.20.432085

**Authors:** Suowen Xu, Sihui Luo, Xueying Zheng, Jianping Weng

## Abstract

Coronavirus disease 2019 (COVID-19) is regarded as an endothelial disease (endothelialitis) with its mechanism being incompletely understood. Emerging evidence has demonstrated that the endothelium represents the Achilles' heel in COVID-19 patients and that endothelial dysfunction precipitates COVID-19 and accompanying multi-organ injuries. Thus, pharmacotherapies targeting endothelial dysfunction have potential to ameliorate COVID-19 and its cardiovascular complications. Primary human umbilical vein endothelial cells (HUVECs) and human pulmonary microvascular endothelial cells (HPMECs) were treated with serum from control subjects or COVID-19 patients. Downstream monocyte adhesion and associated gene/protein expression was evaluated in endothelial cells exposed to COVID-19 patient serum in the presence of KLF2 activator (Atorvastatin) or KLF2 overexpression by an adenoviral vector. Here, we demonstrate that the expression of KLF2 was significantly reduced and monocyte adhesion was increased in endothelial cells treated with COVID-19 patient serum due to elevated levels of pro-adhesive molecules, ICAM1 and VCAM1. IL-1β and TNF-α, two cytokines observed in cytokine release syndrome in COVID-19 patients, decreased KLF2 gene expression. Next-generation RNA-sequencing data showed that atorvastatin treatment leads to a cardiovascular protective transcriptome associated with improved endothelial function (vasodilation, anti-inflammation, antioxidant status, anti-thrombosis/-coagulation, anti-fibrosis and reduced angiogenesis). Treatment of HPMECs with atorvastatin or KLF2 adenovirus ameliorate COVID-19 serum-induced increase in endothelial inflammation and monocyte adhesion by increasing KLF2 expression. Altogether, the present study demonstrates that genetic and pharmacological activation of KLF2 represses COVID-19 associated endothelial dysfunction, heralding a potentially new direction to treat endothelialitis accompanying COVID-19.

## Introduction

Coronavirus disease 2019 (COVID-19) is a severe, yet still ongoing infectious disease caused by a new type of coronavirus-SARS-CoV-2 (1). The pandemic has posed a great threat to public health and imposed tremendous economic burden worldwide. Cardiovascular complications are rapidly emerging as a new threat in COVID-19, and these complications can last even in patients recently recovered from COVID-19, indicate the necessity of assessing the long-term cardiovascular consequences of COVID-19 (2). However, the mechanisms underlying SARS-CoV-2 infection induced dysfunction in the cardiovascular system, remain largely unknown. Since the discovery of viral inclusion structures and inflammatory cell infiltration as well interaction with endothelium in endothelial cells from various vascular beds from patient tissues (3), COVID-19 is considered as an endothelial disease (4), in which multiple aspects of endothelial dysfunction (such as oxidative stress, mitochondrial dysfunction, endothelial cell death, inflammation, glycocalyx disruption, altered cell metabolism etc) occur (5).

The vascular endothelium is not a static bystander, but a metabolically active paracrine, endocrine, and autocrine organ critical for the regulation of vascular tone and homoeostasis, highlighting the possibility that the endothelium represents the Achilles’ heel in COVID-19 patients (6). In particular, in the cascade of injurious responses triggered by cytokine storm accompanying COVID-19, the sentinel vascular endothelium could become dysfunctional, leading to imbalance of tissue homeostasis and ensuing injury. Further elucidation of its pathogenic mechanism is important for developing effective treatments.

In this study, we aim to address the underlying mechanism of endothelial dysfunction in COVID-19 by treating human endothelial cells with serum from COVID-19 patients. We observed that the gene and protein expression of kruppel-like factor 2 (KLF2), a master regulator of vascular homeostasis, was decreased in endothelial cells treated with serum from COVID-19 patients and that genetic or pharmacological activation of KLF2 reverses multiple aspects of endothelial dysfunction. Our study offers a new target for therapeutic intervention of endothelial dysfunction in COVID-19.

## Results

### 1. Patient demographic data

A total of eight severe COVID-19 patients and eight disease-free control patients were used for collecting serum. Demographic data of patients are summarized in Table S1.

### 2. COVID-19 patient serum treated human endothelial cells show KLF2 downregulation, and associated eNOS downregulation, and endothelial inflammation

To test the hypothesis that COVID-19 patients’ serum can cause endothelial dysfunction, we first treated HUVECs with control serum or patient serum for 24 h. Proteins and RNA were collected for western blot and quantitative real-time PCR (qPCR) analysis, respectively. Our data demonstrate that patient serum treatment leads to decreased KLF2 and eNOS protein expression, while increasing ICAM1 and VCAM1 protein expression. qPCR analysis revealed that KLF2, eNOS (also known as NOS3), Thrombomodulin (Thbd) were decreased by patient serum treatment, while VCAM1 gene expression was increased by patient serum treatment (Figure 1).

**Figure 1.**
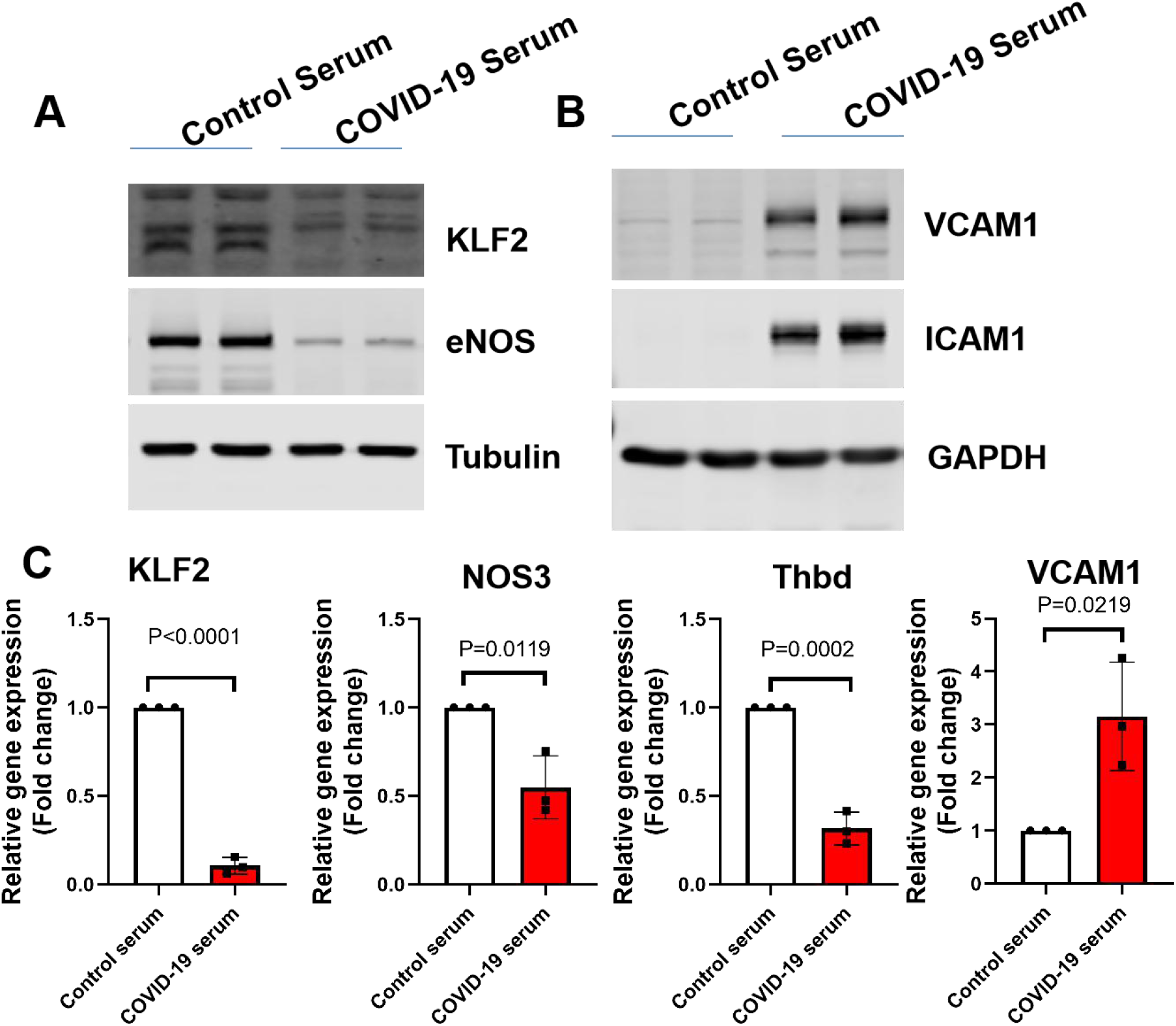
COVID-19 patient serum treated human endothelial cells show KLF2 downregulation, and associated eNOS downregulation, and endothelial inflammation. A. HUVECs were treated with control serum (20%) or COVID-19 serum (20%) for 24 h before protein was collected for western blot analysis of KLF2 and eNOS protein expression. N=3. B. HUVECs were treated with control serum (20%) or COVID-19 serum (20%) for 24 h before protein was collected for western blot analysis of ICAM1 and VCAM1 protein expression. N=3.

### 3. KLF2 is downregulated by components of cytokine storm- TNF-α, and IL-1β

It has been well recognized that COVID-19 is a cytokine release syndrome, in which heightened secretion of TNF-α, IL-1β, IL-6 and many others can trigger inflammatory cell death, local tissue damage as well as systemic multi-organ failure (7). We evaluated whether TNF-α, and IL-1β, two cytokines observed in COVID-19 patients, can decrease KLF2 gene expression. Our data showed that both TNF-α, and IL-1β significantly decreased KLF2 gene expression (Figure 2).

**Figure 2.**
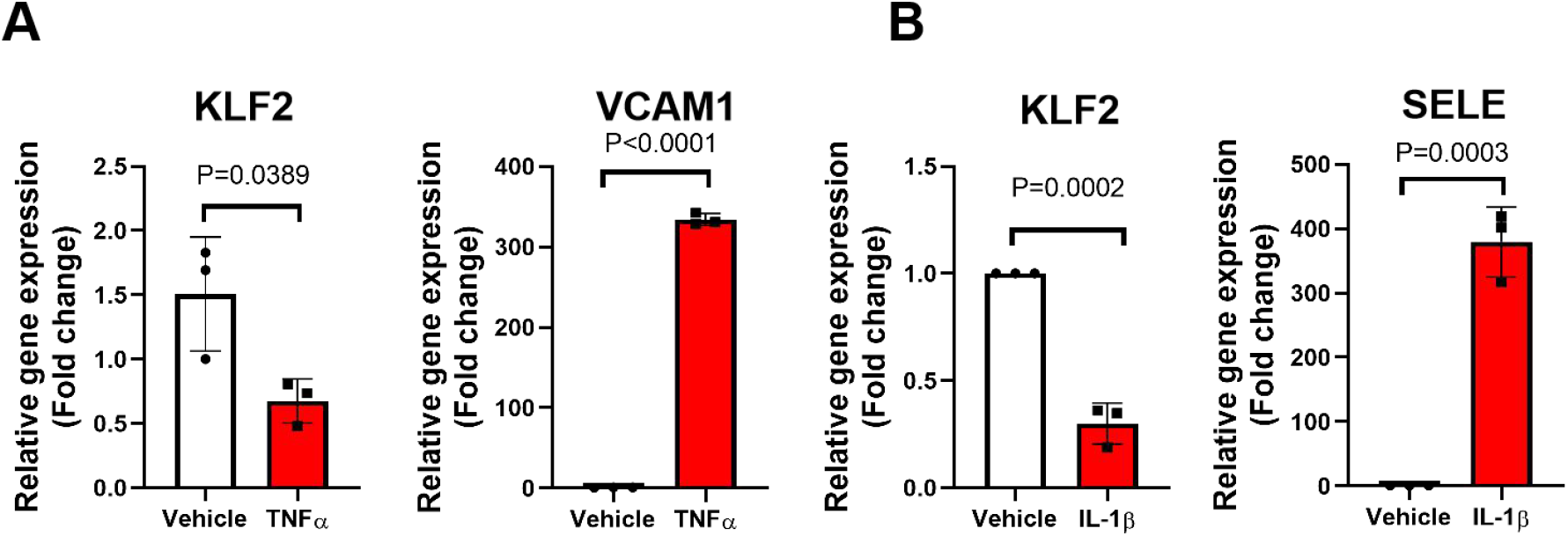
KLF2 is downregulated by components of cytokine storm-TNF-α, and IL-1β. A. HUVECs were treated with vehicle (PBS) or TNF-α (10 ng/ml in PBS) for 6 h before RNA was collected for real-time PCR analysis of KLF2 gene expression using VCAM1 gene as the positive control. N=3. B. HUVECs were treated with vehicle (PBS) or IL-1β (10 ng/ml in PBS) for 6 h before RNA was collected for real-time PCR analysis of KLF2 gene expression using E-selectin (SELE) gene as the positive control. N=3.

### 4. Transcriptional profiling of human endothelial cells treated with atorvastatin in the presence of patient serum

Since statins have reported benefits in cardiovascular outcome in COVID-19 patients (8, 9) and statins are known potent pharmacological KLF2 activators(10, 11). We postulated that KLF2 activation represent one contributing mechanism to COVID-19 associated endothelial dysfunction. We thus performed RNA-sequencing in HUVECs treated with atorvastatin in the presence of patient serum. Our data showed that atorvastatin treatment leads to an overall protective transcriptomic profile (antioxidant, anti-inflammatory, vasodilatory, anti-fibrotic, anti-angiogenesis and anti-thrombotic), which may underscore the potential benefits of statins in COVID-19 treatment (Figure 3).

**Figure 3.**
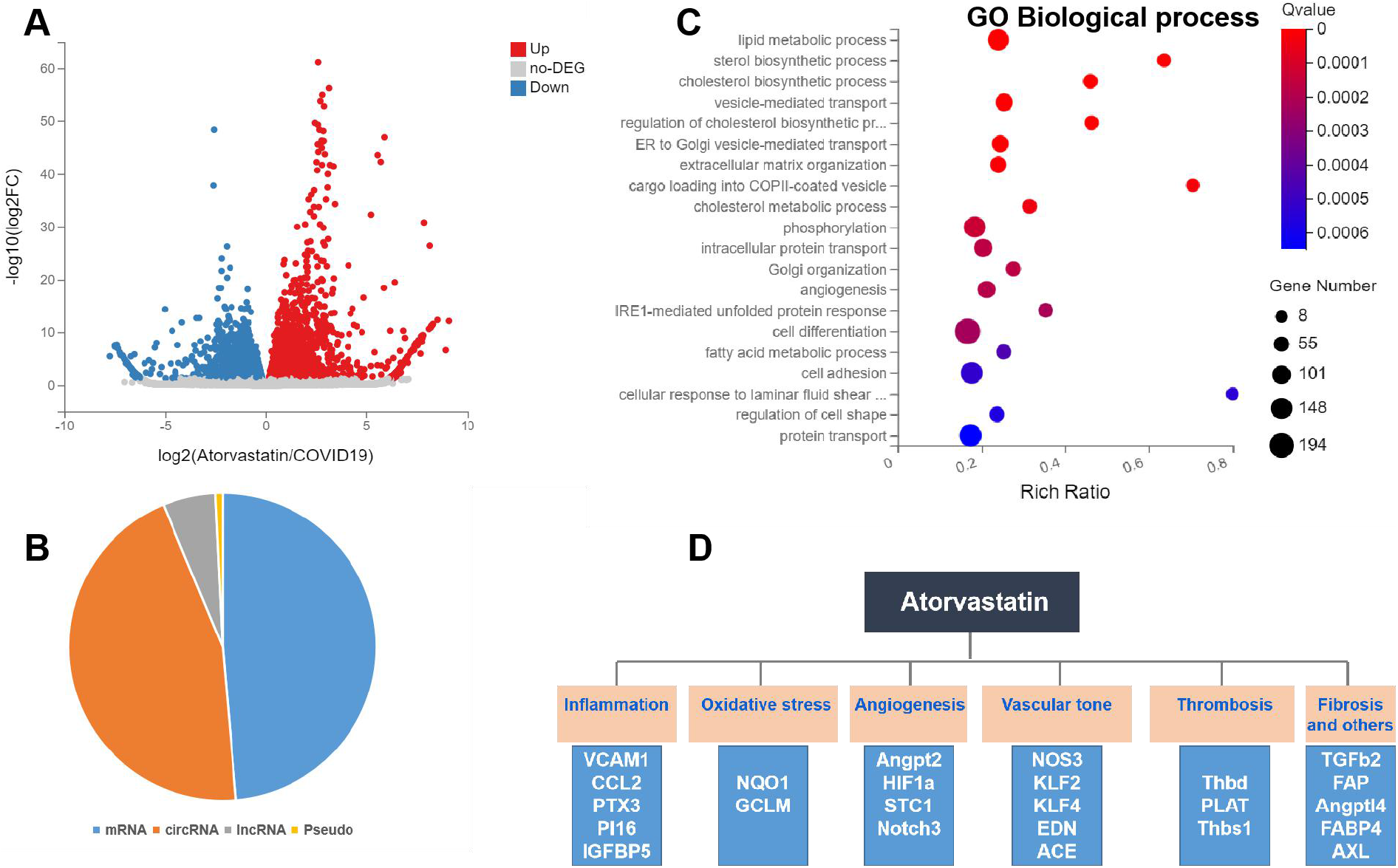
Transcriptional profiling of human endothelial cells treated with atorvastatin in the presence of patient serum. A. Volcano plot of atorvastatin treated HUVECs exposed to COVID-19 patient serum. 4 different donors of HUVECs were treated with vehicle (0.1% DMSO) or Atorvastatin (10 **μ**M in 0.1%DMSO) for 24 h in the presence of COVID-19 patient serum before RNA was collected for next-generation RNA sequencing (RNA-seq). B. Categorization of differentially expressed transcripts in HUVECs treated as described in A. C. Gene ontology (GO) analysis of changes in biological process. D. Summary of differentially expressed genes in response to atorvastatin treatment.

### 5. Atorvastatin regulates the expression of genes relevant to endothelial dysfunction in endothelial cells exposed to patient serum

We further performed qPCR to validate our RNA-sequencing data. Our qPCR data showed that atorvastatin treatment leads to the upregulation of vascular homeostasis related genes (KLF2, KLF4, eNOS, and Thbd), antioxidant genes (NQO1). However, atorvastatin treatment reduces the expression of pro-inflammatory genes (VCAM1, CCL2 and DKK1), vasoconstrictory gene (EDN-1) and anti-angiogenic gene (Angpt2) (Figure 4). These data are of translational significance as increased pulmonary vascular endothelialitis, thrombosis, and angiogenesis are observed in COVID-19 patients (12, 13). In addition, atorvastatin reduces the expression of Angpt2, which emerges a biomarker of endothelial activation in COVID-19 and predicts COVID-19 patients to be admitted to ICU (14).

**Figure 4.**
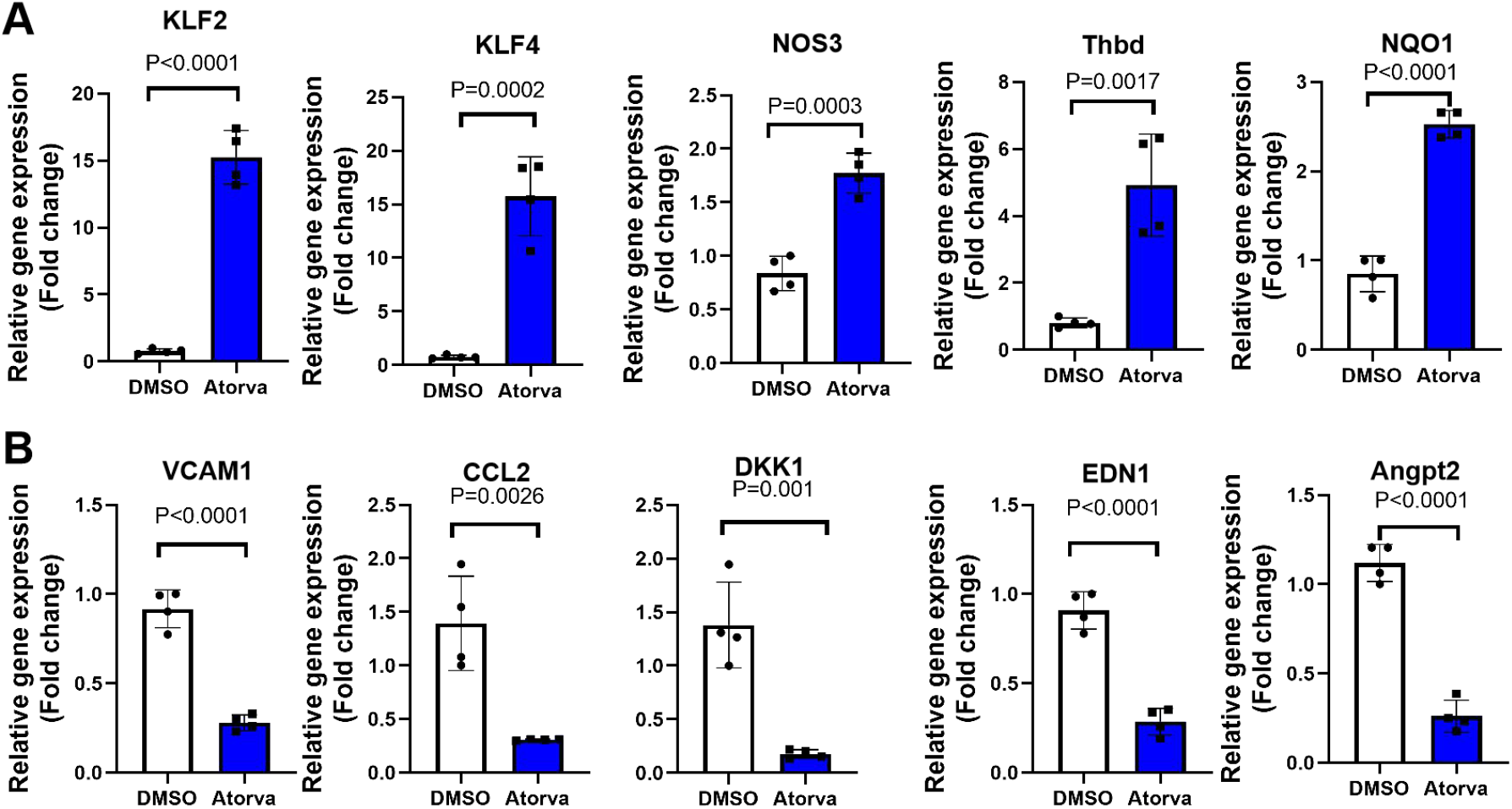
Atorvastatin regulates the expression of genes relevant to endothelial dysfunction in endothelial cells exposed to patient serum. A. HUVECs were treated with vehicle (0.1% DMSO) or Atorvastatin (10 **μ**M in 0.1%DMSO) for 24 h in the presence of COVID-19 patient serum before RNA was collected for real-time PCR analysis of gene expression. Genes related to vascular homeostasis and anti-thrombosis (KLF2, KLF4, NOS3, Thbd) and antioxidant status (NQO1) were presented as fold changes over control. N=4. B. HUVECs were treated as described in A, and expression of genes related to inflammation (VCAM1, CCL2, and DKK1), vascular tone (EDN1 or ET1), and angiogenesis (Angpt2) were presented as fold changes over control. N=4.

### 6. KLF2 overexpression modulates the expression of genes relevant to endothelial dysfunction in endothelial cells exposed to patient serum

Since atorvastatin is a pharmacological activator of KLF2, we next asked whether overexpression of KLF2 via an adenoviral vector can also reverse patient serum induced endothelial dysfunction. Our qPCR data showed that atorvastatin treatment leads to the upregulation of vascular homeostasis related genes (KLF2, eNOS, and Thbd), antioxidant genes (GCLM and NQO1). However, atorvastatin treatment reduces the expression of pro-inflammatory genes (VCAM1, CCL2 and DKK1), vasoconstricotry gene (EDN-1) and anti-angiogenic gene (Angpt2) (Figure 5).

**Figure 5.**
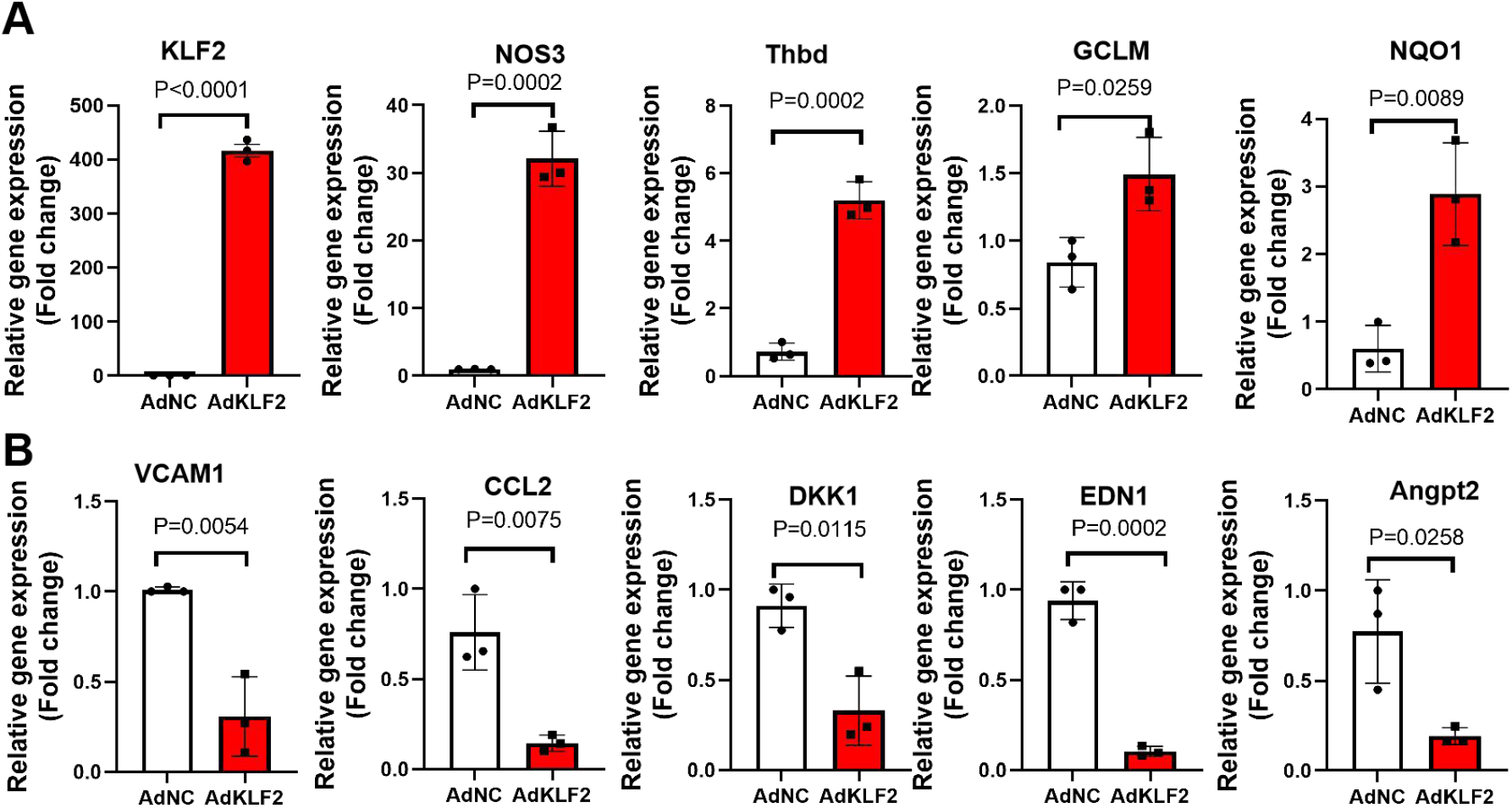
KLF2 overexpression modulates the expression of genes relevant to endothelial dysfunction in endothelial cells exposed to patient serum. A. HUVECs were treated with control adenovirus (AdNC) or KLF2 overexpressing adenovirus (AdKLF2) for 24 h in the presence of COVID-19 patient serum before RNA was collected for real-time PCR analysis of gene expression. Genes related to vascular homeostasis and anti-thrombosis (KLF2, NOS3, Thbd) and antioxidant status (GCLM and NQO1) were presented as fold changes over control. N=3. B. HUVECs were treated as described in A, and expression of genes related to inflammation (VCAM1, CCL2, and DKK1), vascular tone (EDN1 or ET1), and angiogenesis (Angpt2) were presented as fold changes over control. N=3.

### 7. Genetic and pharmacological activation of KLF2 reduced monocyte adhesion to patient serum treated endothelial cells

A recent report has observed significant accumulation of inflammatory cells associated with endothelium, as well as apoptotic bodies (3). In particular, the myocardial arterioles and venules of the COVID-19 patients showed mild to moderate inflammatory infiltrates rich in lympho-monocytic cells (15). However, how these lympho-monocytic cells infiltration in affected tissues/organs was increased is unknown. To this end, genetic and pharmacological activation of KLF2 have been established to reduce monocyte adhesion to activated endothelium (16, 17). We thus investigated the role of KLF2 overexpression and atorvastatin on COVID-19 serum induced monocyte adhesion. Our data demonstrate that both KLF2 overexpression (by KLF2 adenovirus) and activation (by Atorvastatin) significantly reduces monocyte adhesion (Figure 6), providing a proof-of-concept of KLF2 activation in treating endothelial dysfunction in COVID-19.

**Figure 6.**
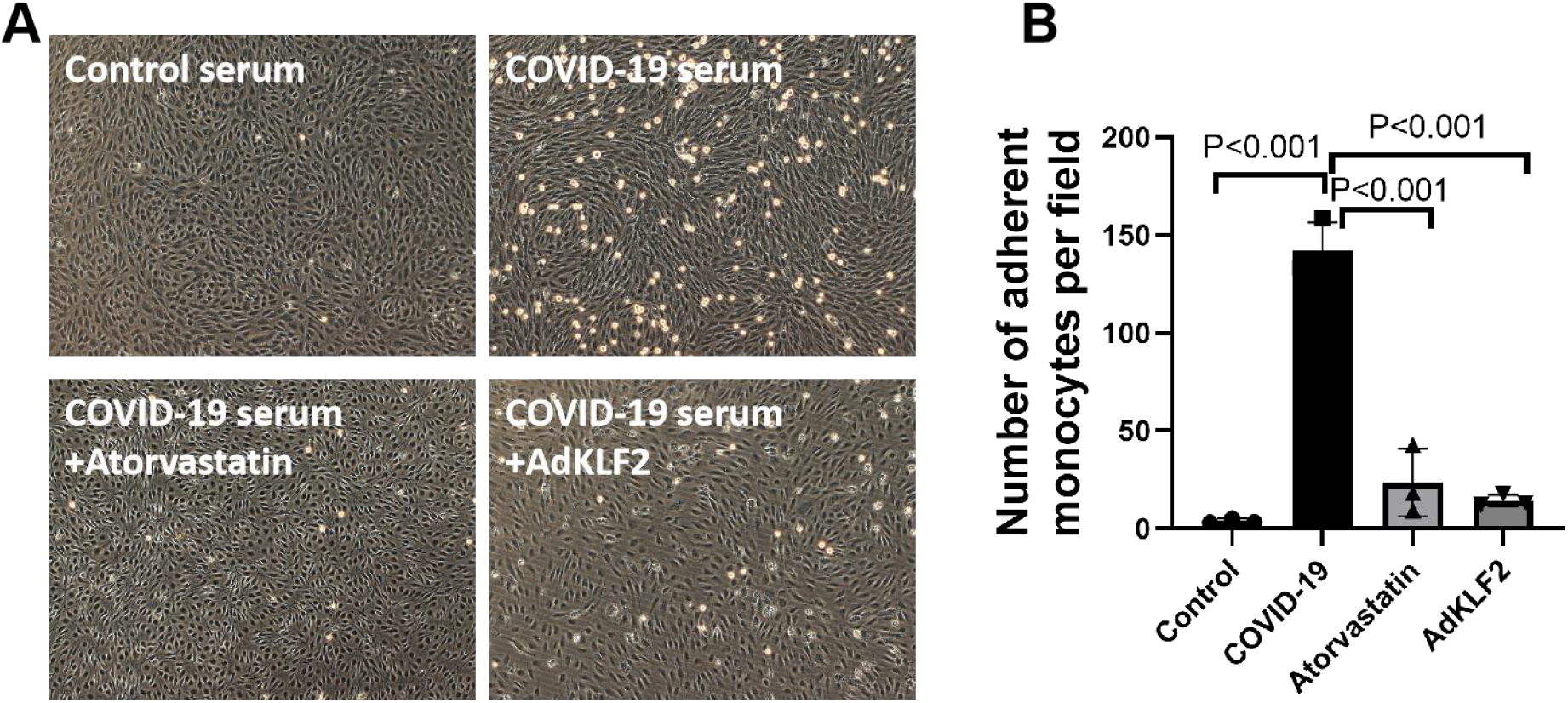
Genetic and pharmacological activation of KLF2 reduced monocyte adhesion to patient serum treated endothelial cells. A. Human pulmonary microvascular endothelial cells (HPMECs) were treated with control serum or COVID-19 serum for 24 h before THP-1 monocyte adhesion assay was performed. Non-adherent cells were washed and photographs of adherent monocytes were taken, N=3. B. A quantification of adherent monocytes as described in A, N=3.

## Discussion

Emerging histopathological evidence from COVID-19 patients have underscored the pivotal role of endothelial cell dysfunction, thrombosis/coagulation, and systemic inflammation in COVID-19 caused by SARS-CoV-2 infection (3, 18). Consecutive inflammatory/immune cell recruitment and endothelial dysfunction could explain microcirculation failure and systemic endotheliitis observed in various vascular beds (15, 19). Mechanistically, elevated levels of TNF-α, interferons, IL-1**β**, IL-6, chemokines (CCL2 or MCP1), and PAI-1 in cytokine storm (20), could relay the chain of endothelial dysfunction, consisting of vasoconstriction, cell injury/death, hyperpermeability, and leukocyte recruitment in the microvasculature. Therapeutic strategies targeting endothelial inflammatory responses and dysfunction could restore the quiescent endothelium and improve multi-organ endotheliitis and injury (21).

### Endothelial dysfunction and endothelialtis in COVID-19

A landmark histopathological study has shown direct SARS-CoV-2 viral infection of endothelial cells evidenced by the presence of viral elements within endothelial cells and the accumulation of inflammatory cells (3). These findings suggest that SARS-CoV-2 infection facilitates endotheliitis in several organs. Since this discovery (3), the importance of endothelial activation and dysfunction in COVID-19 has been intensively pursued (4, 15, 19, 22). More recently, increased markers of vascular inflammation (23), oxidative stress (24), coagulation (12), angiogenesis (12), mitochondrial dysfunction (25), altered endothelial cell metabolism (glycolysis) (25), glycocalyx disruption (26) have been observed in patients with COVID-19 or SARS-CoV-2 spike protein treated endothelial cells (reviewed in (5)). However, the precise mechanism underlying these features of endothelial dysfunction and COVID-19 is largely unknown. Studies focusing on endothelial dysfunction in COVID-19 patients are warranted as to decipher their precise role in severe SARS-CoV-2 infection and multi-organ dysfunction and to identify targets for further interventions (27).

In a recent study, spike protein alone can damage endothelial cells in an AMPK-dependent manner, evidenced by mitochondrial dysfunction, reduced ACE2 expression and eNOS expression and NO bioavailability, and altered endothelial cell metabolism (increased glycolysis) (25). In addition, chloroquine, a disputed anti-COVID-19 medication, may induce endothelial injury/cytotoxicity through lysosomal dysfunction and oxidative stress, the effect of which can be ameliorated by treatment with lysosomal enzyme α-galactosidase A. This suggests that endothelial cell injury may contribute to the failure of chloroquine as effective therapy for COVID-19 partially due to lysosomal dysfunction elicited by chloroquine (28). All these evidences indicate that the endothelium is *bona fide* a key target for devising anti-COVID-19 therapeutics.

However, there are no pharmacotherapies available which specifically target the diseased vascular endothelium in affected organs from COVID-19 patients. Several clinical trials are currently underway to explore this concept. Anti-inflammatory therapies such as colchicine, and tocilizumab (anti-IL-6 receptor monoclonal antibodies) are tested for clinical intervention of endothelial dysfunction in COVID-19 patients (www.clinicaltrial.gov). In our study, we observed that COVID-19 patient serum treated human endothelial cells show features of endothelial dysfunction, including increased monocyte adhesion to activated endothelium, concurrent with decreased expression of vasoprotective molecule KLF2 and eNOS, and increased expression of ICAM1, and VCAM1, raising the possibility that KLF2 serves as a new and promising target for therapeutic intervention of endothelial dysfunction accompanying COVID-19 (Figure S1).

### KLF2 as a negative regulator of endothelial dysfunction in COVID-19

Previous studies have shown that KLF2 is a negative regulator of endothelial activation, dysfunction and thrombosis (29–32). This notion is supported by evidence showing that KLF2 downregulation by treatments with various inflammatory cytokines, such as TNF-α (33) and IL-1β (17), two of which are frequently observed cytokine storm in COVID-19 patients (7). We reproduced the downregulation of KLF2 expression by TNF-α and IL-1β, and further demonstrated that forced overexpression of KLF2 suppressed patient serum induced inflammatory gene and protein (ICAM1 and VCAM1) expression and monocyte adhesion to activated endothelial cells *in vitro*. In addition, KLF2 overexpression reversed patient serum induced eNOS downregulation, indicating that SRAS-CoV2 infection induced cytokine storm disrupts endothelial homeostasis. Therefore, based on our data and published literature (29), it is plausible that KLF2 suppresses COVID-19 induced endothelial dysfunction by multiple mechanisms which may involve: 1) enhancing endothelial quiescence and pulmonary vascular integrity; 2) boosting eNOS dependent NO production; 3) inhibiting ICAM-1, VCAM-1 and E-selectin mediated monocyte adhesion via reported suppressing NF-kB signaling pathway; 4) hemostatic and anti-thrombotic function mediated by thrombomodulin upregulation and PAI-1 downregulation. In our study, we observed that KLF2 was downregulated by COVID-19 patient serum, which disclose a novel mechanism of KLF2 in suppressing endothelial dysfunction in COVID-19. We also observed that adenovirus mediated KLF2 overexpression or atorvastatin reversed COVID-19 patient serum induced monocyte adhesion to endothelial cells, suggesting that pharmacological activation of KLF2 could be a viable strategy of ameliorating endothelial dysfunction in COVID-19.

Based on the protective effects of KLF2 activation, in addition to statins, several drugs or compounds, such as resveratrol (a wine-derived phytochemical) (34), metformin (35), liraglutide (36) have been shown to exert cardiovascular protective actions via KLF2 upregulation. By virtue of the anti-thrombotic and anti-inflammatory properties mediated by KLF2 activation, these drugs/compounds would be expected to lower COVID-19-associated endothelial dysfunction and mortality.

### Mechanism of statin-mediated protective effects against COVID-19

The use of statins was associated with reduced risk for 28-day all-cause mortality, a lower risk of developing severe COVID-19, and faster recovery time (8, 9). However, the protective mechanism of statins in COVID-19 is unclear. We found that statins may be effective by ameliorating endothelial dysfunction triggered by cytokine storm cytokines/chemokines (IL-6, TNF-α, IL-1β, CCL2, interferons etc) released from COVID-19 patients. The possible reasons for statins mediated suppressive effects on endothelial dysfunction are partially due to increased KLF2 expression and regulates the expression of its downstream genes, which includes NF-kB-dependent pro-inflammatory genes (VCAM-1, SELE, CCL2), thrombotic genes (Thrombomodulin, Thromborespondin 1), vascular homeostasis associated genes (eNOS and ET-1 etc). Since statins are pleiotropic drugs with both lipid-lowering and cholesterol-independent effects, both of which may be accountable for the pharmacological effects of statins in COVID-19 patients. In addition, SARS-CoV-2 use ACE2 receptor to achieve entry into host cells, however, the endothelium of the major coronary arteries of COVID-19-positive was devoid of ACE2 receptor expression (15), raising the existence of alternative receptors in endothelial cells. In our study, we do observe that atorvastatin treatment leads to dramatic decrease in the expression of AXL, a new candidate receptor which binds to SARS-CoV-2 spike glycoprotein and facilitates SARS-CoV-2 entry into ACE2-lowly expressed host cells (37), such as the coronary artery endothelial cells. Therefore, the effect of statins on SARS-CoV-2 entry into host endothelial cells is also possible.

### Study limitations

We recognized that the present study has two limitations. First, in light of the presence of SARS-CoV-2 viral inclusion elements within endothelial cells (3), the transcriptomic profile of SARS-CoV-2 infected human microvascular endothelial cells from different vascular beds are warranted to elucidate the gene expression profile after SARS-CoV-2 infection; Second, due to inaccessibility of post-mortem tissues from COVID-19 patients, further detection of KLF2 expression in vascular endothelium from vascular beds from small vessels (such as cardiac capillaries, arterioles and venules) and whether KLF2 expression negatively correlates with increased vascular inflammation, the severity of COVID-19, inflammatory biomarkers and coagulopathy remains to be evaluated.

### Conclusion

The present study uncovers KLF2 downregulation as an important mechanism responsible for COVID-19 induced endothelial dysfunction as well as underscores importance of examining endothelial function in patients recently recovered from COVID-19. From a translational perspective, out study suggests that genetic and pharmacological activation of KLF2 have potential to ameliorate COVID-19 associated endothelial dysfunction, pinpointing a new direction to treat endothelialitis accompanying the devastating pandemic COVID-19.

## Materials and methods

### 1. Patient demographic data

Serum samples were collected from eight confirmed COVID-19 patients as described in our previous cohort (38). All patients were diagnosed with laboratory-confirmed COVID-19 infection and discharged after meeting the National Recovery Standard of COIVD-19 stipulated by the National Health Committee of China. Clinical demographic information and laboratory testing data (Table S1), including age, sex, comorbidities, smoking history, treatment, complete blood counts, blood biochemistry were collected at the time of admission and throughout the course. A subfraction of patients were followed up for various blood tests. Control serum were collected from normal human subjects. This study was conducted under a clinical protocol approved by the Institutional Review Board (IRB) of First Affiliated Hospital of University of Science and Technology of China (protocol number: 2020-XG(H)-009). All participants all agreed to participate in the study and signed informed consents approved by the IRB.

### 2. Drug and adenovirus

Atorvastatin was purchased from Cayman Chemicals (Ann Arbor, MI), and KLF2 overexpression adenovirus with a C-terminal Flag/His tag was custom made at Weizhen Biochem Inc. (Qingdao, Shandong, China).

### 3. Cell culture

HUVECs were isolated from normal pregnant women according to our published protocols with informed consent (39). Three to four different donors of HUVECs were used in this study unless specified otherwise. HUVECs were cultured in ECM media supplemented with 1X endothelial cell growth supplement (ScienCell, Carlsbad, CA), 1X penicillin/streptomycin antibiotic, and 5% FBS. Cells at passage number of 2-4 were used in this study. HPMECs were purchased from ScienCell (Carlsbad, CA) under the identical culture conditions to HUVECs. Endothelial cells were authenticated by staining with endothelial cell marker proteins-CD31 and VE-cadherin as well as DiI-oxLDL uptake.

### 4. Real-time quantitative PCR (qRT-PCR)

Total RNA was extracted from cultured human ECs using a RNeasy Mini kit (Qiagen). For reverse transcription, total RNA was converted into first strand complementary DNA (cDNA) using a Reverse Transcription Kit (Takara) following the manufacturer’s instructions. Quantitative real-time PCR was then performed with a Roche LC96 Real-Time PCR Detection System, using SYBR Green Supermix (Roche) for relative mRNA quantification. The sequences of all the primers used were listed in Table S2.

### 5. Western blot analysis

Whole cell lysates were prepared from cultured cells. For Western blots (39), total cell lysates (15-20 μg) were separated by SDS-PAGE, transferred to nitrocellulose membrane (Pall, East Hills, NY) and were subsequently blocked in LI-COR blocking buffer (LI-COR Biosciences, Lincoln, NE) at room temperature for 1 h. Then, the blots were incubated overnight at 4 °C with appropriate primary antibodies listed in Table S3. Then after being washed 3 times with 1 X Tris buffered saline with 0.1% Tween-20 (TBST), membranes were incubated with IRDye^®^ 680RD Goat anti-Mouse IgG (H+L) or IRDye^®^ 800CW Goat anti-Rabbit IgG (H + L) (1:10,000 dilution in 1XTBST; LI-COR) at room temperature for 30 min. Images were visualized by using an LiCor-CLX Infrared Imaging System (LI-COR).

### 6. Assay of monocyte adhesion to endothelial cells

Human THP-1 monocyte adhesion assay was performed as we previously described (39, 40). In brief, HUVECs were incubated with 20% patient serum (combined from 2 patients with 1:1 ratio) or control serum for 24 h before addition of THP-1 monocytic cells (1.5X10^4^) for 30 min. Non-adherent cells were washed and images were taken. Three images at different optic fields were taken for assessing the number of adherent monocytes.

### 7. Next-generation RNA-sequencing

HUVECs were treated with atorvastatin (10 uM) for 24 h before treatment with COVID-19 patient serum for 24 h. After that, RNA was extracted using the RNA Minieasy Kit (Qiagen). The sequencing data was filtered with SOAPnuke (v1.5.2) by (1) Removing reads containing sequencing adapter; (2) Removing reads whose low-quality base ratio (base quality less than or equal to 5) is more than 20%; (3) Removing reads whose unknown base (‘N’ base) ratio is more than 5%, afterwards clean reads were obtained and stored in FASTQ format. The clean reads were mapped to the reference genome using HISAT2 (v2.0.4). Bowtie2 (v2.2.5) [3] was applied to align the clean reads to the reference coding gene set, then expression level of gene was calculated by RSEM (v1.2.12). The heatmap was drawn by pheatmap (v1.0.8) according to the gene expression in different samples. Essentially, differential expression analysis was performed using the DESeq2 (v1.4.5) with Q value ≤ 0.05. To take insight to the change of phenotype, GO (http://www.geneontology.org/) and KEGG (https://www.kegg.jp/) enrichment analysis of annotated different expressed gene was performed by Phyper (https://en.wikipedia.org/wiki/Hypergeometric_distribution) based on Hypergeometric test. The significant levels of terms and pathways were corrected by Q value with a rigorous threshold (Q value ≤ 0.05) by Bonferroni.

### 8. Statistical analysis

Data are presented as means ± SD unless otherwise indicated. Statistical analysis was performed using GraphPad Prism Software Version 8.3 (GraphPad software, La Jolla, CA). Results were evaluated by t-test or by one-way analysis of variance (ANOVA) when appropriate. When multiple comparisons were made, a Bonferroni correction was performed for each test. A P value less than 0.05 were considered to be statistically significant.

## SUPPLEMENTARY DATA

Supplementary data are available online.

## ACKNOWLEDGEMENTS

We thank Siqi wang and Haitao Wang for technical supports; Xiumei Wu and Yujie Liu for assisting human umbilical cord collection and culture of HUVECs.

## AUTHOR CONTRIBUTIONS

J.W. and S.X. conceived the project and guided the study. S.X. performed experiments, statistical analysis and wrote the manuscript. S.L. and X.Z. collected clinical samples from patients and provide insightful discussions. J.W. supervised the study. All the authors have read, revised and approved the manuscript.

