## Supplemental data for "KLF2 is a therapeutic target for COVID-19 induced endothelial dysfunction"

### Table S1 Patient demographic data

| ID | AGE (yr) | WBC (10E9/L) | Lymphocyte (%) | ALT (U/L) | AST (U/L) | hs-CRP (mg/L) | BUN (mmol/L) | Sample collection since onset (day) |
|---|---|---|---|---|---|---|---|---|
| XY1900358 | 70 | 4 | 17.6 | 33 | 20 | 7.4 | 3.74 | 17 |
| XY1900410 | 80 | 8.9 | 5 | 101 | 48 | 25.2 | 8.88 | 14 |
| XY1900429 | 55 | 10.54 | 16.7 | 18 | 13 | 20.7 | 9.25 | 18 |
| XY1900434 | 64 | 9.75 | 6.8 | 41 | 31 | 24.5 | NA | 19 |
| XY1900437 | 59 | 10.38 | 12.6 | 229 | 94 | 12.3 | 9.86 | 18 |
| XY1900503 | 93 | 9.84 | 22.1 | 34 | 37 | NA | 8.33 | 17 |
| XY1900509 | 70 | 5.48 | 23.4 | 72 | 40 | 3.9 | 11.72 | 13 |
| XY1900510 | 67 | 7.01 | 21.8 | NA | NA | 0.5 | NA | 29 |

**Table S2 Primer list**

| Gene name | Forward | Reward |
|---|---|---|
| KLF2 | CACGCACACAGGTGAGAA | ACAGATGGCACTGGAATGG |
| KLF4 | GAACCCACACAGGTGAGAAA | GTAGTGCCTGGTCAGTTCATC |
| NOS3 (eNOS) | CCGGAACAGCACAAGAGTTA | GTCTGTGTTACTGGACTCCTTC |
| VCAM 1 | GGCTTGTGTGTTCGGTTTC | GGAGCTCTACTCATTCCCTAGA |
| CCL2 (MCP1) | GTCCCAAAGAAGCTGTGATCT | AGTCTTCGGAGTTTGGGTTTG |
| GCLM | GAGTTGCACAGCTGGATTCT | CCTCCCAGTAAGGCTGTAAAT G |
| NQO1 | GGGATGAGACACCACTGTATT T | TCTCCTCATCCTGTACCTCTTT |
| EDN1 | AAGGCAACAGACCGTGAAA | GTCTTCAGCCCTGAGTTCTTT |
| Angpt2 | GTGACTGCCACGGTGAATAA | GGGTCCTTAGCTGAGTTTGATG |
| Thbd | ACGTGGATGACTGCATACTG | ACCAGGTCGTAGTTAGGGTAG |
| DKK1 | AGCACCTTGGATGGGTATTC | CTGATGACCGGAGACAAACA |
| SELE | GTGTATGTCCTCTGGAGAATGG | GAACCCATTGGCTGGATTTG |
| GAPDH | GATTCCACCCATGGCAAATTC | CTGGAAGATGGTGATGGGATT |

**Table S3 Source of antibodies used**

| Antibody used | Vendor | Catlog No. |
|---|---|---|
| KLF2 | ZenBio Inc, Chengdu, China | 863587 |
| eNOS | ZenBio Inc, Chengdu, China | 250094 |
| ICAM1 | ZenBio Inc, Chengdu, China | 384648 |
| VCAM1 | ZenBio Inc, Chengdu, China | 220417 |
| GAPDH | ProteinTech | 60004-1-Ig |
| Tubulin | ProteinTech | 11224-1-AP |

**Figure S1**

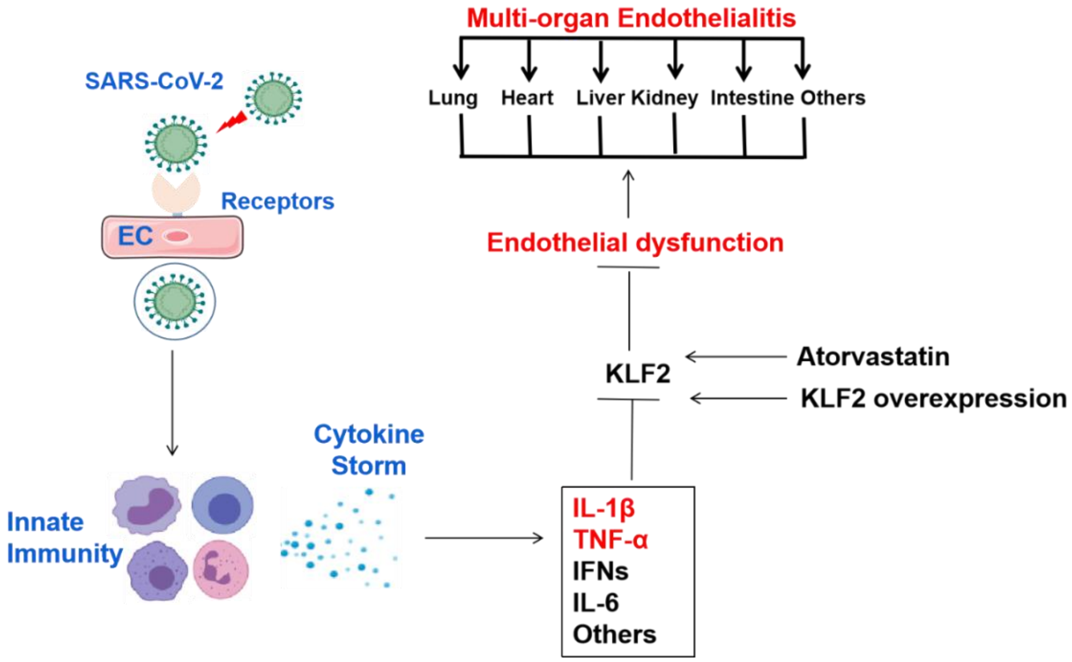


**Endothelial dysfunction contributes to COVID-19-associated multi-organ endothelialtis: potential role of KLF2**

The role of endothelial dysfunction in COVID-19 has gained intensive research interest. Growing evidence suggests that the angiotensin converting enzyme 2 receptor (ACE2 receptor) is expressed on endothelial cells (ECs) from small vessels (such as capillaries) in the lung, heart, kidney, and intestine. Upon viral infection of ECs or neighboring cells by severe acute respiratory syndrome coronarvirus 2 (SARS-CoV-2), ECs become activated and dysfunctional. SARS-CoV-2 infection in endothelial cells is believed to trigger a cytokine storm that plays a critical role in the pathogenesis of endothelialitis and vascular injury, eventually leading to KLF2 downregulation， NF-kB activation, vascular dysfunctions and respiratory as well as multi-organ failure in COVID-19 patients. As a result of endothelial activation and endothelial dysfunction (ED), the levels of pro-inflammatory cytokines (IL-1, IL-6, and TNF-α), chemokines (MCP-1), and acute phase reactants (IL-6, CRP, and D-dimer) are elevated. Therefore, ED contributes to COVID-19-associated vascular inflammation, particularly endotheliitis, coagulopathy in the lung, heart, kidney, intestine. Further research of the important role of ED in COVID-19 patients is warranted to offer targets for therapeutic intervention.
